# emObject: domain specific data abstraction for spatial omics

**DOI:** 10.1101/2023.06.07.543950

**Authors:** Ethan A. G. Baker, Meng-Yao Huang, Amy Lam, Maha K. Rahim, Matthew F. Bieniosek, Bobby Wang, Nancy R. Zhang, Aaron T. Mayer, Alexandro E. Trevino

## Abstract

Recent advances in high-parameter spatial biology have yielded a rapidly growing new class of biological data, allowing researchers to more comprehensively characterize cellular state and morphology in native tissue context. However, spatial biology lacks a cohesive data abstraction on which to build novel computational tools and algorithms, making it difficult to fully leverage these emergent data. Here, we present emObject, a domain-specific data abstraction for spatial biology data and experiments. We demonstrate the simplicity, flexibility, and extensibility of emObject for a range of spatial omics data types, including the analysis of Visium, MIBI, and CODEX data, as well as for integrated spatial multiomic experiments. The development of emObject is an essential step towards building a unified data science ecosystem for spatial biology and accelerating the pace of scientific discovery.

## Main

Tissue is a key level of biological organization where diverse collections of cells work together to perform vital functions. The composition and spatial configuration of cells within the tissue are precisely tuned, and even small perturbations in spatial structure can result in dysregulation of tissue function. Disruption of the spatial organization within tissue is implicated in numerous disease processes and can be highly specific.^1^.

While the study of tissue structure in disease and in clinical diagnostics has long been investigated via hematoxylin and eosin (H&E) staining and immunohistochemistry (IHC), recent technological advances have enabled highly-multiplexed detection of proteins and RNAs in their native tissue context. Techniques for *in situ* molecular measurements (“spatial omics”), including an array of methods for spatial transcriptomics and antibody-based spatial proteomics, build upon a rich history of histopathological analysis ^2–14^. These new methods enable the study of spatial patterns and aid in mechanistic inference with newfound molecular richness. Recent novel insights from these approaches include the discovery of cellular neighborhoods associated with patient survival in colorectal cancer, tissue morphological features associated with ductal carcinoma progression, and cell microenvironments that predict patient prognosis in head and neck cancers ^15–17^.

Spatial omics data is fundamentally cross-modal; spatial assays often jointly capture both molecular and imaging data: for instance Visium (10X Genomics) captures transcriptomic data via next generation sequencing as well as a H&E reference image. This requires that analytical workflows also be cross-modal; for example, researchers might seek to extract the molecular features (e.g. gene counts or protein level) from a spatial region that corresponds to a region of interest observed in H&E data (**Figure 1A**). Moreover, structured or unstructured information is often associated with entire samples in a spatial experiment (e.g. metadata that describes sample disease state), spatial ROIs (e.g. morphological tissue landmarks), or single cells (e.g. cellular morphology features). Thus, a robust spatial analysis requires fluid movement between images, spatial ROIs (“masks”), and annotations assigned to various spatial scales (**Figure 1A-B**).

**Figure 1:**
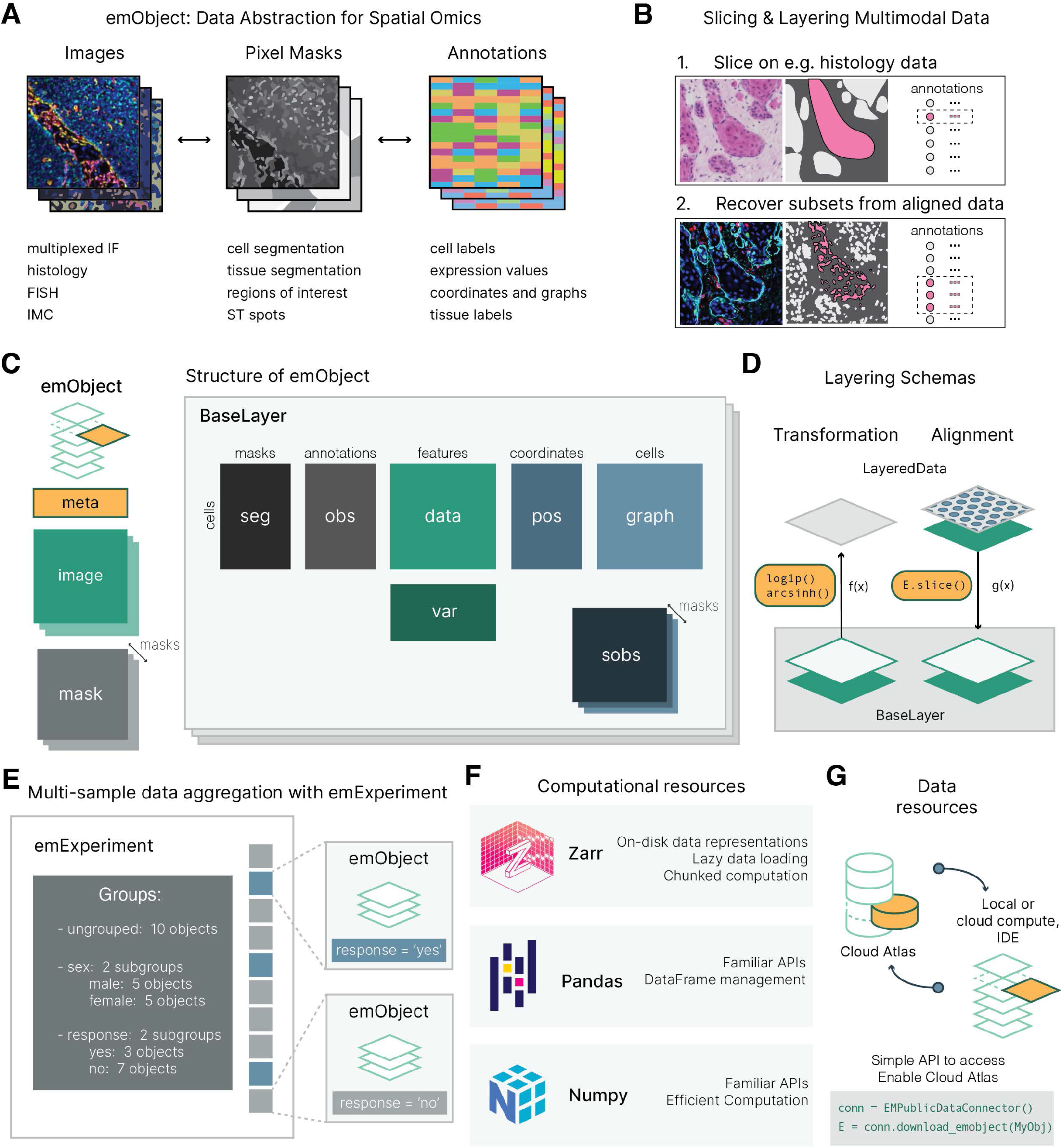
emObject is a domain specific data abstraction for spatial omics. **(a)** Spatial omics datasets are multimodal and multiscale. Spatial omics acquisitions are composed of images, pixel masks (e.g. ROI annotations), and annotations, which can include both structured (e.g. cell by marker expression) and unstructured (e.g. sample level metadata) data. **(b)** emObject facilitates seamless transitions between operations on images, masks, and annotations. For example, emObject can store multimodal spatial data in “layers’’ and facilitates “slicing” of data on the basis of mask segments (Panel 1) to retrieve spatially-defined subsets of the entire emObject (Panel 2). **(c)** emObject contains dedicated attributes for all aspects of a spatial data acquisition. emObject’s core data attributes are organized into layers, which contain the central data matrix, *data*, the spatial coordinates, *pos*, annotations on observations (e.g. cell types), *obs*, mapping of observations to mask segment, *seg*, annotations on variables, *var*, and graphs derived from spatial information, *graph.* Core data attributes are aligned along shared observation and variable axes where possible. emObject contains additional attributes for images, masks, and metadata. **(d)** emObjects are organized into layers. Layers can contain either transformations on data (left) or hold aligned multimodal spatial data acquisitions (right). **(e)** emExperiment facilitates the analysis of cohorts of emObjects. emExperiment can aggregate sample level metadata, assemble cohorts on the basis of metadata covariates, and apply analysis over groups of emObjects. **(f)** emObject is built on popular open-source libraries from the Python data science ecosystem including Zarr, Pandas, and NumPy. **(g)** emObject is designed to be integrated with cloud computing infrastructure to provide streamlined access to data atlases.

As spatial omics data generation has accelerated, an impressive array of statistical and computational methods have been developed to address the novel data science questions raised by this emergent data class ^18^. However, in practice, the overall usability of these methods is limited, in part because there are no clear standards for representing spatial omics data in a unified and consistent format. Domain specific data abstractions are not new to biology; in molecular biology, AnnData ^19^ provides a consistent, domain-specific approach to representing single cell RNA sequencing (scRNA-seq) data, and in population genetics, Hail’s MatrixTable provides a representation of GWAS data suitable for the distributed cloud computing required to perform biobank-scale linear algebra. In both cases, data abstractions have provided more accessible frameworks for working with complex data, and become important tools in their respective fields. However, to date, no such standard exists in spatial omics, and Thus, a robust, flexible solution to unlock seamless and transferable spatial omics analytical workflows is urgently needed.

To this end, we introduce emObject, a domain-specific data abstraction that is purpose-built for the intricacies of spatial omics data. emObject enables seamless translations between the core elements of a spatial omics dataset - images, masks, and annotations. We extend the notion of shared indexing used in other libraries to apply to spatial attributes at a variety of scales (**Figure 1C**). Importantly, emObject is data type agnostic and is well-suited for both spatial transcriptomics and spatial proteomics, and can efficiently represent all data types associated with a single spatial assay. Additionally, we introduce a lightweight wrapper on emObject, emExperiment, to facilitate the management of multiple spatial data acquisitions and perform cohort-based analysis. Recognizing the richness of the existing Python data science ecosystem, we designed emObject with a familiar API to users of existing tools. This choice increases usability, but also makes emObject workflows portable, and able to connect to libraries that power downstream analysis, such as PyTorch for machine learning workflows.

## Results

emObject conforms to the existing notion of aligning attributes of spatial experiments along common data axes. As with previous data abstractions, like AnnData, emObject is organized around a core data matrix, ‘.datà, which is indexed by cells and contains cell-wise entries of values for each variable (i.e. genes in spatial transcriptomics or proteins in spatial proteomics). Annotations on cells or variables are stored in separate attributes, ‘.obs’ and ‘.var’, which are indexed by cells and variables, respectively. Because emObject is purpose-built for spatial omics data, we provide a dedicated attribute, ‘.pos’, to store spatial coordinates of observations (**Figure 1C**).

Many spatial annotations are performed on masks, discrete pixel arrays that indicate a label over some spatial region of interest. To accommodate this class of annotation, emObject extends the concept of aligned data axes to a new axis: segments. We define a segment as a single continuous region within a mask. For example, a mask of two disconnected tumor regions in a tissue would contain two segments. We allow annotations on segments to be stored in a new data object ‘.sobs’. ‘.sobs’ itself is aligned with the data matrix via the ‘.seg’ attribute, which is indexed by observations, and maps observations to segments to preserve a consistent link between observations at varying spatial scales. Notably, though emObject is centered on the notion of observations, which for most workflows will be cells or spots, nothing about the design precludes the analysis of acellular data. In fact, emObject facilitates this via the ability to define annotations and analysis on any image segment.

emObject implements a notion of *layering* to accommodate both transformations on data (e.g. holding both raw and normalized data of the same modality in a single object) and aligned multimodal datasets (e.g. storing aligned Visium + CODEX samples from adjacent tissue sections in the same object) (**Figure 1D**). Additionally, we also define functionality for integrated analysis of multimodal spatial datasets in multilayered emObjects that contain aligned data. For instance, we define a *slice* operation that uses associated segmentation masks to extract corresponding regions of aligned datasets.

A single emObject represents all of the data collected from a single spatial data acquisition on a single sample; this ensures the link between all attributes of the data abstraction. However, most spatial experiments contain multiple samples. To accommodate multiple acquisitions, we developed a superclass for managing multiple emObjects, emExperiment. emExperiment provides a single container for multiple emObjects and provides tools for cohort analysis. Because many spatial experiments rely on differential analysis across groups, emExperiment allows the user to build groups of emObjects on the basis of metadata covariates associated with an emExperiment’s constituent emObjects, and apply custom analysis functions over these groups, facilitating the automation of cohort analysis (**Figure 1E**).

Throughout emObject and emExperiment, we leverage the rich Python data science ecosystem to provide a familiar API to the user. As many spatial omics assays generate imaging data, either as the primary data (e.g. multiplexed proteomic or transcriptomic imaging) or as accessory data (e.g. 10X Visium), we implemented efficient image representation using Zarr, an open-source standard for high-dimensional array data^20^. Our image container allows large image data to reside on-disk, rather than in-memory, and for only relevant chunks of large images to be loaded into memory when needed. Additionally, because Zarr implements efficient compression, our container reduces disk requirements for imaging data. In certain assays where image channels correspond with biomarkers or genes, images can also be aligned along a shared variable axis. Other emObject attributes are built using NumPy and Pandas, giving emObject attributes a familiar API ^21–23^. We also note that this approach prepares emObject for interaction with analysis tools that require parallel processing, for example via Dask (www.dask.org) integrations with emObject’s Zarr backend (**Figure 1F**).

Finally, emObject is designed to be integrated with cloud computing environments and petabyte scale data repositories. We present an early vision for a broader ecosystem that facilitates a seamless flow between data access, data analysis, method development, and cloud computing. To that end, we have generated emExperiments for 4 publicly available datasets, comprising 265 Zarr files total, which can be reproducibly queried and loaded into emObject (**Figure 1G**). Additionally, we provide example analysis workflows that interact with emObject as interactive notebooks that accompany this initial release. These workflows illustrate the natural multimodal analysis capabilities of emObject through simple yet intuitive examples, and can easily be extended to encompass rapid advancements in spatial assay development.

To highlight the utility of emObject, we performed a few exemplar analyses on various spatial datasets including 10X Visium, H&E, NanoString GeoMX, CODEX, and MIBI. We use these examples to demonstrate two integrated analyses - one between H&E and spatial transcriptomics and a second between spatial transcriptomics and proteomics. As spatial data modalities expand and become more commonplace in the omics data stack, integrative multiomic analysis of these datasets provides a unique opportunity to capture a more complete view of how the transcriptome maps to proteomic or histological observables and vice versa.

First, to demonstrate emObject’s ability to interact with multimodal data and flexibly move between features derived from different components of a spatial dataset (**Figure 1A**), we performed an analysis of publicly available 10X Visium data of the mouse olfactory bulb. Although typically framed as a spatial transcriptomics assay, 10X Visium data is multimodal, as it contains both the spatiomolecular information and a backing H&E stain. These two data types are aligned by the 10X processing pipeline. We clustered the Visium spots by gene expression, which reveals the layered spatial organization of the olfactory bulb (**Figure 2A**).

**Figure 2:**
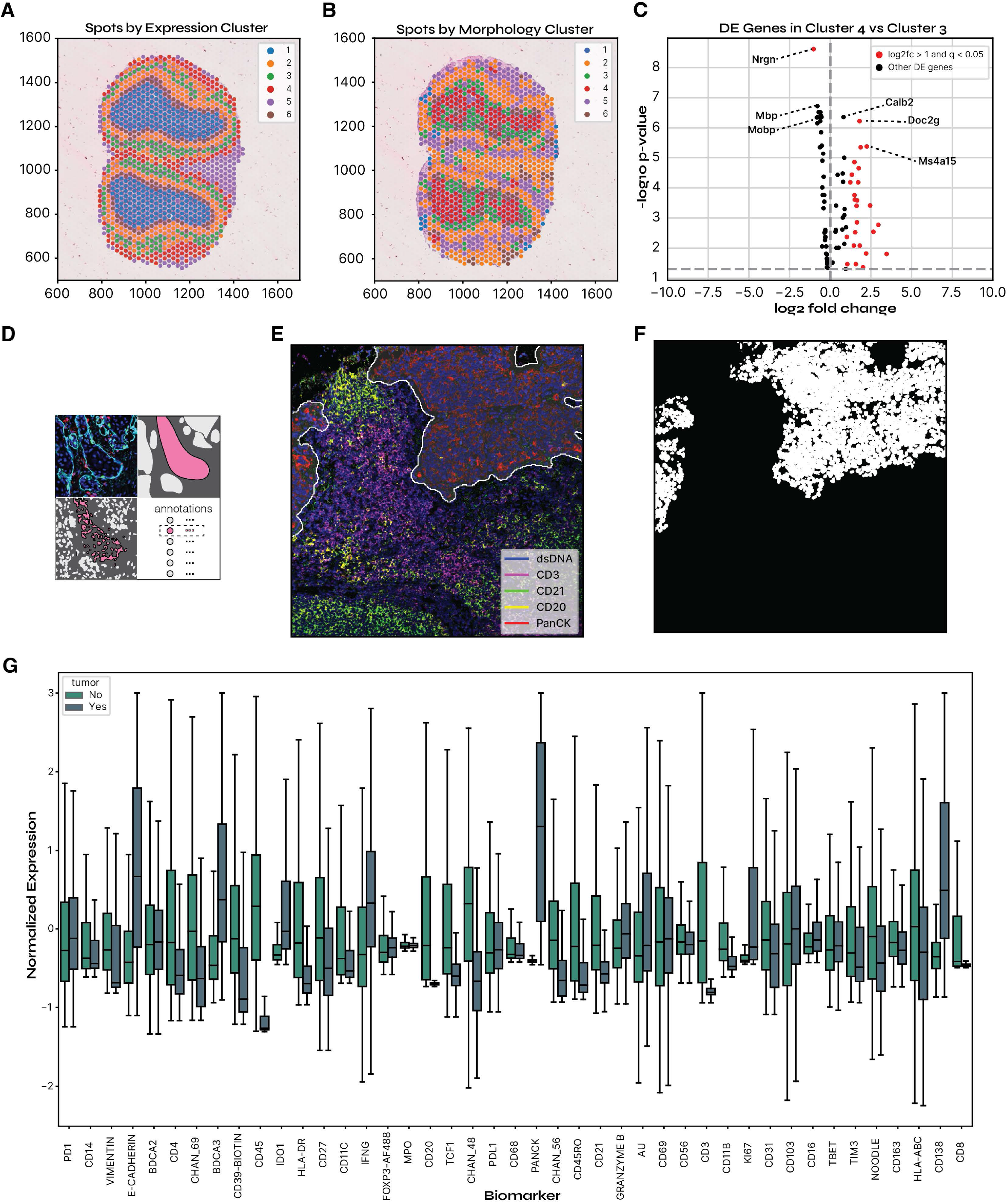
emObject facilitates a variety of spatial omics analysis workflows. **(a)** Spatial transcriptomics of the mouse olfactory bulb. Visium spots are overlaid on the associated H&E stain of the mouse olfactory bulb. Spots are colored by gene expression clusters. **(b)** Morphology clusters of the mouse olfactory bulb. Visium spots are overlaid on the associated H&E stain of the mouse olfactory bulb. Spots are colored by their morphology cluster, as determined by the H&E image analysis. **(c)** Differential expression of genes between two morphological clusters (red and green in *b*). Volcano plot shows differentially expressed genes (points) arranged according log2 fold change (log2fc) (x) and -log10 corrected p-value of a Wilcoxon rank sum test (y). Points in red have both an adjusted p-value < 0.05 and log2fc>1. Vertical dashed line indicates log2fc=0. Horizontal dashed line indicates adjusted p-value of 0.05. **(d)** emObject unlocks seamless workflows between images masks, and, annotations. Starting from an image, users can define masks (top right), extract corresponding observations (bottom left), and their associated annotations (bottom right). **(e)** Example visualization of a mask managed by emObject. Mask segment associated with tumor regions is highlighted with white overlay and bounded by a white line. A subset of multiplexed image channels are shown, including dsDNA (blue), CD3 (magenta), CD21 (green), CD20 (yellow), and PanCK (red). **(e)** Example visualization of extracting corresponding cells for a mask segment. Cell segmentation mask (white) is shown for the subset of cells found within the mask segment highlighted in *(d)*, as computed by emObject. **(f)** Segment-based analysis of marker expression. Boxplot illustrates the normalized expression (y-axis) of each biomarker (x-axis) for cells within or outside of the mask segment highlighted in *(d)* (colors). Cells in the tumor region are shown in blue; cells outside the tumor region are shown in green.

Because recent works have shown that rich molecular information can be predicted from histopathological data alone, we were interested in whether the H&E data conveyed the same spatial information as the gene expression clusters ^24, 25^. We divided the backing H&E image into 32 x 32 px tiles, featurized image tiles using a deep neural network, and clustered the resulting feature vectors into the same number of clusters as for gene expression (**Figure 2B, *Methods***). We observed that the segmentation of the olfactory bulb into regions on the basis of morphology was similar to the segmentation on the basis of gene expression (**Figure 2A**). We further observed that the most interior compartment as defined by the gene expression clusters (i.e. blue region in **Figure 2A**) was split into primarily 2 clusters in the morphology example (i.e. red and green clusters in **Figure 2B**). We used emObject’s segment observation and mask handling functionality to extract the corresponding Visium spots for each morphological cluster (***Methods***). We used this information to identify differentially expressed genes between the two compartments (**Figure 2C**). We found that several dendrite associated genes (*Nrgn, Mbp, Mobp*) had decreased expression in morphology cluster 3 relative to morphology cluster 4. We also found several calcium signaling associated genes to be more highly expressed in morphology cluster 3 relative to morphology cluster 4, including *Calb2*, *Doc2g*, and *Ms4a15*. The spatial patterning of Visium spots of each morphology cluster suggests differential localization of soma and dendritic spines in the interior of the mouse olfactory bulb. Thus, an integrated analysis of H&E and spatial gene expression provides clues as to the possible origin of differences in spatial features.

emObject’s mask management capabilities are flexible to different data modalities and accommodate cell segmentation masks, transcriptomic spots, and masks that represent regions of interest (ROIs). To illustrate, we performed a simple ROI based analysis on the MIBI dataset associated with *Rahim et.al. 2023* to compare biomarker expression on the basis of ROI membership (***Figure 2D***). We constructed an emObject of an example data acquisition from this study, and annotated the tumor region of the tissue using a ROI mask (**Figure 2E**). We extracted the corresponding cells for the tumor region (**Figure 2F**) and non-tumor region. Finally, we compared biomarker expression values between the tumor and non-tumor region, observing that certain markers such as PanCK, E-Cadherin, and CD138 were more highly expressed in the tumor region versus non-tumor region. While this is a straightforward analysis, emObject substantially streamlines the exchange of information between images, masks, and annotations, reducing this workflow to just a few lines of code.

We further extended emObject’s mask and ROI functionalities to aligned multimodal datasets (**Figure 1B, 1D, 3A**). Multimodal spatial datasets are increasingly common and provide a more comprehensive representation of the spatial-molecular processes at play in tissue. Here, we use data collected on nearby tissue sections in breast cancer to construct a multimodal emObject that contains two layers: one containing a 10X Visium data and the other containing CODEX data. We spatially aligned the two datasets via an affine transformation computed between the backing H&E stain of the Visium data and the DAPI channel of the CODEX data (**Figure 3B-C**). Using emObject’s *slice* function and layer-specific segmentation masks, we spatially subsetted the CODEX layer using the location and size of Visium spots in the aligned coordinate system, returning only CODEX cells that fall within the boundaries of an aligned Visium spot (**Figure 3D, *Methods***). We observe that, of spots that align with CODEX cells, Visium spots contain between 1 and 12 cells, consistent with the value reported by 10X Genomics (**Supplementary Figure 1**) (10X Genomics). In the subsetted CODEX data, we performed a standard cell expression clustering to identify cell types (**Supplementary Figure 2A)**^26^. We then performed neighborhood analysis on the CODEX-phenotyped cells to identify patterns in the composition of localized cell groups (**Supplementary Figure 2B)**. Initial analysis yielded six neighborhood types (NTs): two NTs (NT2 and NT3) were composed primarily of PanCK+ tumor cells, two stromal neighborhoods (NT1 and NT4), one aSMA+ Pdpn+ stromal neighborhood (NT5), and one CD68+ enriched neighborhood (NT0) (**Supplementary Figure 2C**). We selected NT5 to examine whether cells in the aSMA+ Pdpn+ stromal neighborhood existed in similar or distinct transcriptional states (**Figure 3E-F**). Next, we used the aligned Visium gene expression layer to examine whether NT5 existed in distinct gene expression states (**Figure 3G, *Methods***). We found that nearly half of all cells in NT5 existed in Visium Expression State 0 (V0), with a sizable subpopulation in Visium Expression State 6 (V6) (**Figure 3H**). We investigated marker genes for V0 and V6 (**Supplemental Table 1**), and found that V0 contained marker genes associated with tissue remodeling, cell adhesion, and ESR positivity, while V6 contains marker genes associated with immune response. Both V0 and V6 contained marker genes that were indicative stromal fibroblasts, including TMSB4X (V6) and POSTN (V0), which was consistent with the observed morphology in NT5. This suggests that similarly composed cellular neighborhoods as defined by proteomics, like NT5, can exist in distinct transcriptional states, which could not be detected by a unimodal analysis.

**Figure 3:**
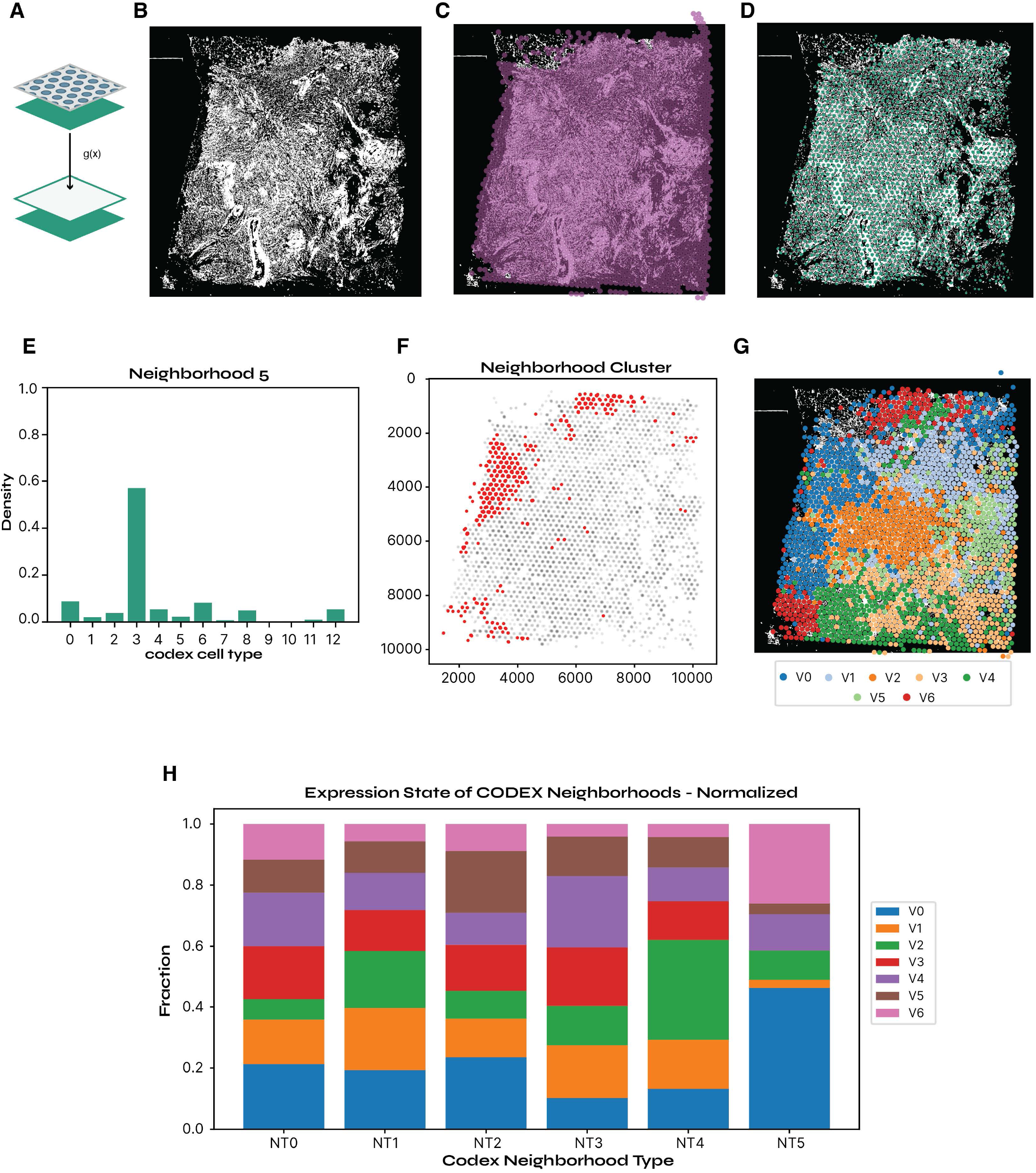
emObject accommodates aligned multimodal spatial datasets. **(a)** Slicing aligned datasets with emObject. We define a spatially-aware querying strategy, slicing, that extracts corresponding spatial observations from aligned datasets. **(b)** CODEX sample of breast cancer tissue. Cell segmentation mask is colored white where a cell is present and black otherwise. **(c)** Visium and CODEX alignment. Same image as in **(c),** with aligned Visium spot locations overlayed in magenta. **(d)** CODEX cells contained within aligned Visium spots. Results of emObject slice operation to return the corresponding CODEX cells of the Visium spots in **(c).** Same image as in **(b)**, but overlaid with a point at each cell centroid of a cell retained after the slice operation (green points). **(e)** Composition of Neighborhood Type 5 (NT5). Bar chart illustrates the probability density (y-axis) of each of the cell types (x-axis) in NT5. **(f)** Spatial distribution of CODEX cells featurized as members of NT5. Plot shows the centroid of each CODEX cell retained after the slice operation (gray points). Cells featurized as members of NT5 are highlighted (red points). **(g)** Aligned spatial location of the Visium expression clusters. Same image as **(b)** overlaid with Visium spots that contain at least one CODEX cell are colored by their expression cluster. **(h)** Expression state of CODEX neighborhoods. For each CODEX-defined neighborhood type (x-axis), bar chart shows the proportion of cells in that neighborhood (y-axis) that are within the bounds of Visium spots of each expression cluster (colors). Colors correspond to **(g)**.

emObject also provides simple tooling for working with publicly available data, facilitated by Enable Atlas’s public data repositories. We generated emObjects from MIBI data as presented in *Rahim et. al. 2023,* which is available as a public study on the Enable Atlas (**Figure 4B**) ^27^. We grouped the MIBI data using a single metadata covariate, treatment status, resulting in two cohorts: surgery (SOC) and atezolizumab (immunotherapy) treatment. We then defined an experiment level analysis workflow to compute Ki-67 positivity in cells, via emObject annotation (**Figure 4C**). Finally, we analyzed the spatial structure of Ki-67+ cells, comparing the two cohorts. By comparing the Ripley’s K between images from each cohort, we observe a mild left shift in the K curve in patients receiving immunotherapy, indicating increased aggregation of Ki-67+ cells (e.g. left shift from 500-1500px radius), consistent with the study’s finding of Ki-67 enrichment in T cell neighborhoods ^27^. However, the effect is small and further analysis would be required to make definitive conclusions about the spatial organization of Ki-67+ cells (**Figure 4D**). Nonetheless, quantitative analysis of tissue organization is uniquely enabled by spatial omics, and cohort comparison is facilitated by emObject and emExperiment.

**Figure 4:**
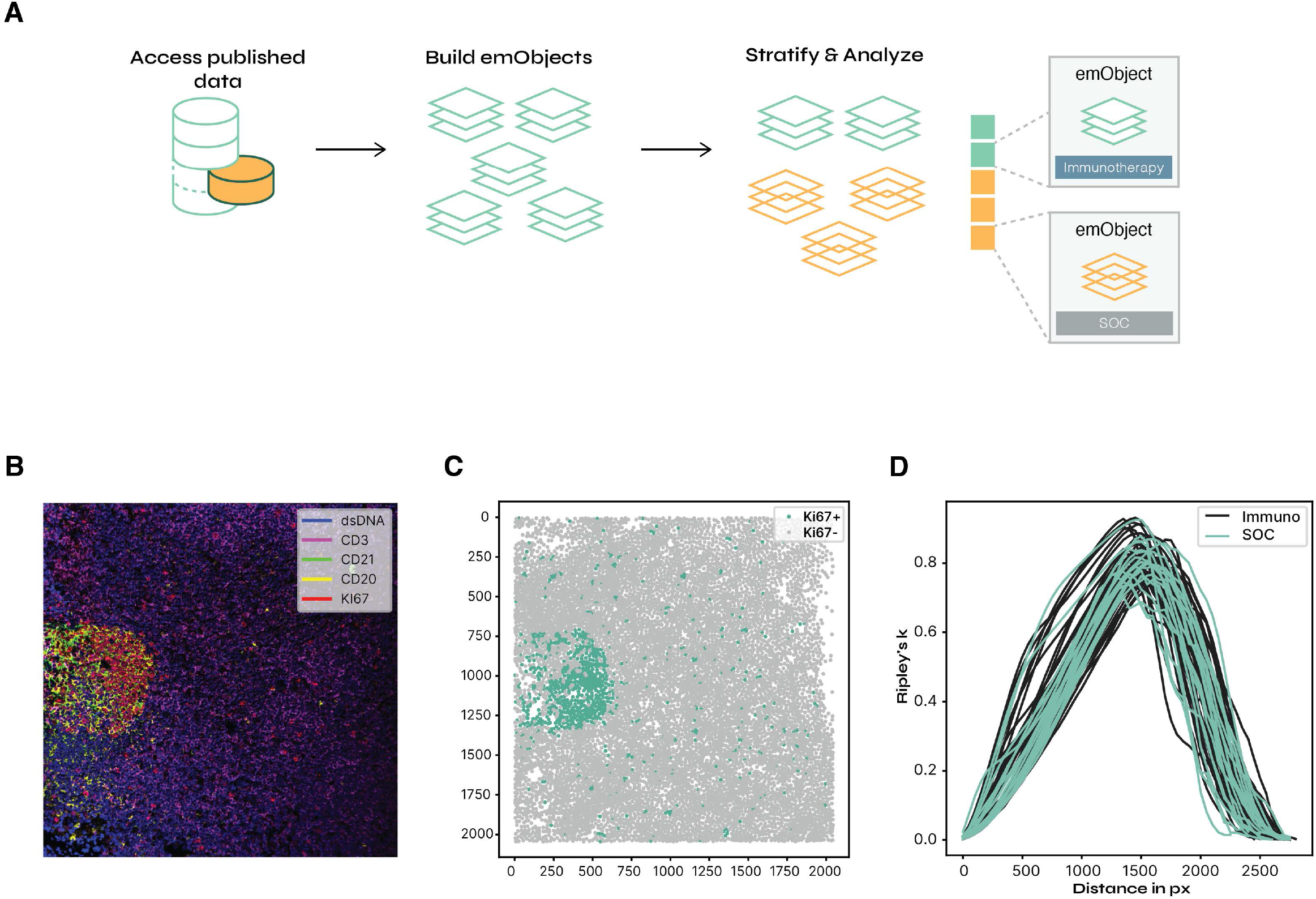
emExperiment manages analysis across multiple emObjects. **(a)** Suggested workflow for emExperiment. Atlas data is accessed and assembled into multiple emObjects, which are managed by emExperiment. emExperiment facilitates cohort assembly on the basis of a metadata covariate, and custom defined analysis workflows can be applied to analysis cohorts. **(b)** Sample MIBI image from *Rahim* et. al. 2023. **(c)** Spatial distribution of Ki67+ cells (green) among all cells (grey) present in the image presented in **(b)**. **(d)** Comparison of Ripley’s K curves for two analysis cohorts - patients receiving immunotherapy (black) or standard of care (green). Curves show the estimate of Ripley’s K (y-axis) as a function of distance in pixels (x-axis).

To demonstrate the extensibility of emObject and emExperiment, we developed a package to compute human interpretable cellular morphology features and integrated this into a workflow for cell phenotyping with both morphological features and surface protein expression. We created an emExperiment representation of five CODEX images of head and neck cancer samples. We used emExperiment to iterate through each sample, extracting morphology features from whole-cell segmentation masks. We then organized the newly extracted spatial features with their respective image in the emExperiment data representation. To complete cell phenotyping, we aggregated morphology and surface marker expression from all images in the emExperiment. We then performed an unsupervised analysis on the integrated cellular information to uncover the signatures of major immune cells and tumor cells in the samples, and wrote these cell type annotations back to the emExperiment (**Figure 5B**). We identified two types of tumor cells in the sample: PanCK+Siglec8+ tumor cells and PanCK+Siglec8-tumor cells. The PanCK+Siglec8+ cell population highly expresses PanCK, Ki67, and Siglec8. Morphologically, these cells are small sized and highly circular. We also found that the cells have irregular boundaries from their low solidity measurement. On the contrary, the PanCK+Siglec8-population expresses lower PanCK and Ki67. This cell population has large cell areas and are highly eccentric compared to the PanCK+Siglec8+ tumor cell population. In the immune cell branch, B cells are found to be low in extent and solidity. Further in-depth analyses can be performed to identify precise cell types, conclude cellular interactions, and construct prediction models for clinical outcomes (**Figure 5B**). emObject facilitated the development of the morphology package by providing consistent, organized representations of the data, allowing the scientific developer to focus on answering research questions of interest rather than on data management.

**Figure 5:**
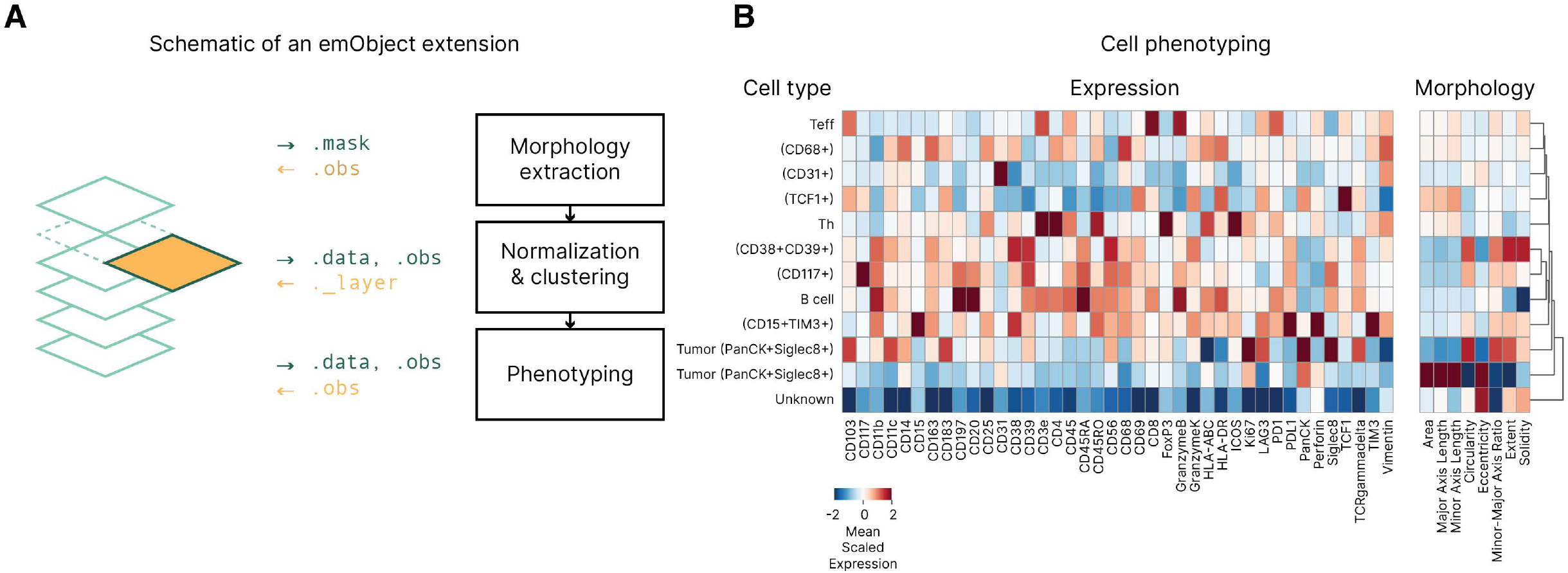
emObject extensibility. **(a)** Morphology analysis extension built on emObject. The extension extracts cellular morphology from *.mask* and writes morphology features to *.obs*. Then it performs normalization & clustering with data from *.data* and *.obs* and writes back to *._layer*. Finally, using information from *.data* and *.obs*, the cells were phenotyped and the annotations are written back to *.obs*. **(b)** Cell phenotyping using the combination of cellular protein marker expression and morphology. Scaled expression and morphology are shown, clipped at [-2, 2]. Cell populations were arranged based on hierarchical clustering.

## Discussion

Here, we introduced emObject, a domain specific data abstraction for spatial omics. Furthermore, we demonstrated the wide-ranging functionality of emObject for assorted spatial analysis workflows, including working with masks and ROIs, subsetting data with spatial information, performing analysis of aligned multimodal spatial omics dataset, building tools on top of emObject, and organizing larger datasets into analysis cohorts with emExperiment.

A key objective we had for defining a data abstraction for spatial omics data was to streamline the future development of new analytical tools as part of a data science ecosystem. emObject fulfills this objective by providing an expected common input to downstream analysis libraries. emObject provides dedicated attributes for annotations at all spatial scales and on all components of a spatial dataset, providing an intuitive workflow for analysis tools to interact with emObject. At a high level, this flow consists of a tool reading relevant emObject data, which is stored in an expected location and format given the emObject schema, performing computations in the analysis package, and writing results back to an emObject as new annotations (**Figure 5A**). In this way, emObject unifies results of distinct analyses and aggregates them in a single object.

Importantly, we view emObject as a single piece of a unified spatial omics analysis ecosystem. With emObject, we tackled the problem of defining a data abstraction and API for interaction with complex spatial datasets; however, this is just a single component of what researchers do with spatial data. For example, data visualization and interaction with gigapixel images is a substantial challenge in spatial omics and digital pathology. Additionally, given the large size of spatial omics datasets, simple storage, management, and access of large quantities of spatial omics data is a significant, and costly, challenge. Similarly, these properties of spatial omics datasets present new problems for efficient and cost-aware computation, particularly in deep learning applications. Moreover, as spatial omics datasets become increasingly common, more researchers, including those without a bioinformatics background, are becoming interested in spatial datasets, generating a need for simple, low or no-code analysis tools. Finally, we recognize that a truly unified biological data science ecosystem must also encompass non-spatial assays, including connections to other formats and analysis toolkits. While some solutions have been proposed to address these challenges, the current state of tooling and computational resources in the field is largely piecemeal. We believe that a complete analysis ecosystem would allow seamless integration between each of the aforementioned components, and that emObject represents an important first step in that direction.

Our experience with the development of emObject has underscored the importance of thinking about biological data analysis workflows with a wide perspective and eye towards integration. We believe that the highest quality software tools will abstract analysis workflows, for example multimodal spatial analysis, rather than overfitting to assay-specific intricacies, and integrate across the biological data stack.

## Methods

### Mouse olfactory bulb expression and morphology

The Visium dataset used for analysis of the mouse olfactory bulb was downloaded from 10X Genomics (publicly available on the 10X Genomics website). To compute morphological features, we extracted the high resolution H&E image from the dataset and manually annotated the region of the image that contained tissue. Using this annotation, we generated non-overlapping tiles of 32x32px from the image. We excluded tiles that contained less than 80% tissue area from downstream analysis.

To compute morphological clusters on Visium tiles, we featurized image tiles using a pretrained ResNet-152 neural network. The resultant feature vectors were clustered using k-means clustering as implemented in scikit-learn v. 1.2.2 ^28^. We set k=6, as this matched the number of clusters present in the provided gene expression clusters of the original dataset.

We then generated a pixel segmentation mask to map regions of the H&E image to their corresponding image tile (e.g. the area of the mask corresponding to the image tile was filled with an integer identifier that corresponds to the tile). We stored this mask in an emObject, and stored the morphological cluster label information in .sobs.

Gene expression information was provided with the dataset, and we used these cluster labels in our downstream analysis of image clusters. To perform differential expression analysis between morphology clusters that mapped to the same expression cluster, we extracted the corresponding expression of Visium spots that were within the bounds of image patches assigned to the morphology cluster of interest. We concatenated the spot expression for all patches of the same type and performed a differential expression analysis using a Wilcoxon Rank Sum test, as implemented in diffexpy v0.7.3 (https://github.com/theislab/diffxpy).

### Aligned CODEX + Visium Analysis

#### Data acquisition

We obtained near-adjacent tissue sections from a breast tumor from Dr. Nancy R. Zhang (University of Pennsylvania). CODEX data were generated by Enable Medicine using a standard 51-plex marker panel. Visium data was generated by Biochain (Newark, CA).

#### CODEX data preprocessing

We performed cell segmentation using DeepCell v. 0.12.2 with the Mesmer algorithm as implemented in the Enable Portal at https://app.enablemedicine.com/portal/visualizer ^29^.

#### Image registration

We aligned the CODEX and Visium data by computing an affine transformation between the high resolution H&E image and the DAPI channel of the CODEX image using a custom in-house pipeline. We used the affine transformation matrix to project the coordinates of Visium spots into the coordinate system of the CODEX data, completing the alignment process.

#### Defining Visium segmentation mask

In order to use emObject’s slice functionality, we defined a segmentation mask that corresponded to the Visium spots. Similar to standard format for cell segmentation masks, the Visium segmentation mask is an integer matrix that is 0 everywhere except for the locations of Visium spots. We mapped spot barcodes to a unique integer identifier. We then filled all matrix entries within the boundary of a Visium spot with the corresponding unique identifier. To compute the boundary of each Visium spot, we extracted the spot size and scale factor from the Visium metadata, and centered a circle of diameter *spot size * scale factor* at the corresponding spot centroid.

#### Slicing multimodal emObjects

To find CODEX cells that were contained within Visium spots, we used emObject’s slice function on a multimodal emObject in which one layer contained the aligned Visium data and the other contained CODEX data. We used the Visium layer as the “anchor layer” and the CODEX layer as the “target layer” in emobject.slice, which computes the intersection of segmentation masks and uses the unique segment IDs in target layer to subset the entire emObject. For all mask-based queries, we retain all “target layer” cells that have any amount of overlap with the “anchor layer”.

#### Defining CODEX cell types

To define CODEX cell type information, we computed marker intensities within each segmented cell that was retained after the slice information. Raw marker intensities were normalized using an arcsinh transform followed by marker-wise z-scoring. Normalized marker intensities were clipped to the interval [-3, 3]. We used these transformed marker intensities to compute a k-NN graph in expression space (k=15). We then partitioned the graph using Leiden clustering as implemented in leidenalg v 0.9.1 to obtain 12 cell clusters used in this analysis ^30^. Clusters were manually interpreted to determine cell type identity.

#### Defining gene expression states of Visium data

To determine gene expression clusters of the aligned Visium spots, we first performed a standard preprocessing pipeline to our Visium data. We filtered out all spots that had total counts outside the interval [1000, 35000] or had >20% mitochondrial gene counts. We also filtered out genes that were expressed in fewer than 10 spots. Data were normalized by total counts by cell and log1p transformed. Lastly, the data were subsetted to the top 2000 most highly variable genes. We performed dimensionality reduction with PCA (50 components), followed by the construction of a k-NN graph (k=15), and Leiden clustering to determine spot clusters. All Visium preprocessing and clustering was performed using scanpy v. 1.9.3 ^31^.

#### Determining the expression state of CODEX neighborhoods

We defined the expression state of a CODEX neighborhood by the gene expression of Visium spots corresponding to aligned CODEX neighborhoods. To find the CODEX neighborhoods contained within specific Visium spots, we again used emObject’s slicing functionality, this time one Visium spot ID at a time, to map each Visium spot to its corresponding CODEX cells. Because each CODEX cell was assigned to a neighborhood type, this allows us to compute, for each CODEX neighborhood type, the distribution of Visium spot expression states in which that neighborhood resides.

### Cohort analysis with emExperiment

#### Data access and cohort assembly

We obtained the data from *Rahim* et. al. 2023 via the Enable Atlas and manually constructed emObjects for each MIBI data acquisition^27^. For the purposes of this analysis, we retained only cell expression values and cellular coordinates in the emObjects. We used the provided metadata to define two analysis cohorts on the basis of treatment status - patients were either received immunotherapy or standard of care. We applied our downstream analysis to these cohorts.

#### Defining Ki67+ Cells

We obtained established thresholds for marker positivity from the study authors. For this analysis, we defined Ki67 expression intensities above 0.733 as Ki67+ and all other cells as Ki67-.

#### Computing Ripley’s K statistic

We computed the Ripley’s K statistic as:

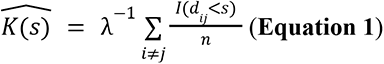

where *λ* is the average point density, *I* is an indicator function returning the count of events of interest within distance *s* and *n* is the total number of possible events. Because Ripley’s K is estimated at single distances, we generated a K curve by computing K at every 50 px distance increment, bounded by the overall image dimension. Because we were interested in the spatial co-occurrence of Ki67 cells, we defined the “event of interest” as the detection of a Ki67 cell, and computed this distribution over each Ki67 cell as a possible starting point for the distance.

### Morphology analysis extension

#### CODEX image segmentation

We performed cell segmentation using DeepCell v. 0.12.2 with the Mesmer algorithm whole cell segmentation as implemented in the Enable Portal at https://app.enablemedicine.com/portal/visualizer ^29^.

#### Extracting cellular protein expression

We computed the cellular protein expression from marker intensities within each segmented cell as implemented in the Enable Portal at https://app.enablemedicine.com/portal/visualizer.

#### Extracting cellular morphology features

We extracted cellular morphology features from cell segmentation masks. The morphology features are defined as follows.

| Morphology Feature | Description |
| --- | --- |
| **Area** | Number of pixels occupied by the cell mask. |
| **Major axis length** | Major axis length from the ellipse with the same normalized second central moments as the cell mask. |
| **Minor axis length** | Minor axis length from the ellipse with the same normalized second central moments as the cell mask. |
| **Circularity** | Calculated as $4 * \pi * area / perimeter^2$, where perimeter is the length around the cell mask measured through the centers of peripheral pixels that are 4-connected (pixels that share an edge with the central pixel). |
| **Eccentricity** | The ratio of the distance between focal points to the major axis length of the ellipse with the same normalized second central moments as the cell mask. |
| **Minor-major axis ratio** | The ratio of minor axis length to major axis length. |
| **Extent** | The ratio of area to the number of pixels in the bounding box of the cell mask, formulated as *area / (bounding box rows \* bounding box columns)*. |
| **Solidity** | The ratio of area to the number of pixels in the convex hull image, which is the smallest convex polygon that contains the cell mask. |

The second central moment *μ_2_* of a cell mask or an ellipse is computed as:

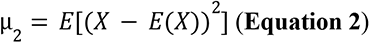

where *X* is a *n * 2* matrix composed of the *n* points in the region and *E* is the expectation

operator. Area, major axis length, minor axis length, eccentricity, extent, solidity, and perimeter were extracted using scikit-image v0.20.0.

#### Defining cell types

To define the cell types in five CODEX images, we combined two cell feature types, protein expressions and morphology measurements, from all images. Raw protein marker intensities were normalized using an arcsinh transform followed by marker-wise z-scoring. Raw morphology measurements were normalized by morphology feature-wise z-scoring. Normalized marker intensities and morphology measurements were clipped to the interval [-2, 2]. We constructed a k-NN graph (k=10) in the integrated space of these transformed marker intensities and morphology measurements. We then partitioned the graph with Leiden clustering using leidenalg v0.9.1 to obtain cell clusters ^30^. The clusters were manually interpreted to determine cell type identity. Clusters of the same cell type were merged, and 12 cell type populations were identified.

## Code availability

The core functionality of emObject is available as an open-source Python package at https://gitlab.com/enable-medicine-public/emobject. We welcome community contributions to the codebase. We also include documentation and sample notebooks at the same link.

## Data availability

The Visium mouse olfactory bulb dataset is freely available from 10X Genomics at https://www.10xgenomics.com/resources/datasets/adult-mouse-olfactory-bulb-1-standard-1. The MIBI dataset was previously published in *Rahim* et.al. 2023 under doi: 10.17632/2zgppyr2rr.1 ^27^. Data used for morphology-aware phenotyping is previously published in *Wu* et. al. 2022 and is available under the terms of that publication^17^. Aligned CODEX+Visium dataset is available via Enable Atlas and via emObject’s public data access library. Additionally, 275 emObjects from 4 public studies are available via emObject’s public data access library (see documentation at https://gitlab.com/enable-medicine-public/emobject/-/blob/main/notebooks/AccessingPublicDataDemo.ipynb) or via the Enable Atlas Portal at https://app.enablemedicine.com/portal/visualizer. After creating a free academic account, researchers can access data used in this publication and other public datasets via the Portal Data Manager.

## Author contributions

E.A.G.B, A.E.T, and A.T.M conceived the study. E.A.G.B implemented emObject with assistance and advice from A.E.T. M.H. contributed morphology analysis and code. A.L. constructed publicly available datasets as emObjects for release with this manuscript. M.F.B. developed and performed Visium+CODEX image registration. N.R.Z provided tissue used in Visium+CODEX analysis. M.K.R. provided annotations, insights, and assistance with the dataset from *Rahim et. al 2023*. B.W. provided technical advice and assistance with the implementation of emObject. E.A.G.B, A.E.T, M.H., A.L. and A.T.M wrote the manuscript. All authors reviewed the final manuscript.

---

**Supplementary Figure 1:**
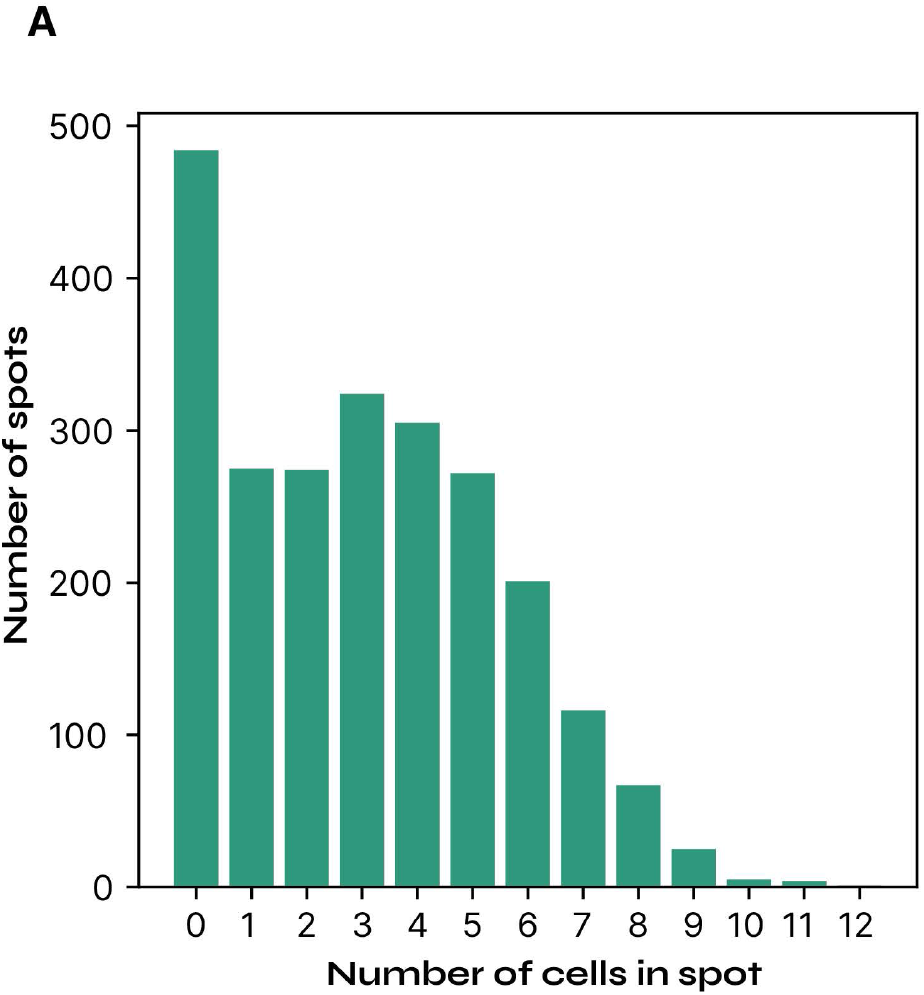
Counts of aligned CODEX cells in Visium spots. **(a)** Bar chart shows the number of Visium spots (y-axis) that contained a specific count of CODEX cells (x-axis) as computed by the emObject slice operation.

**Supplementary Figure 2:**
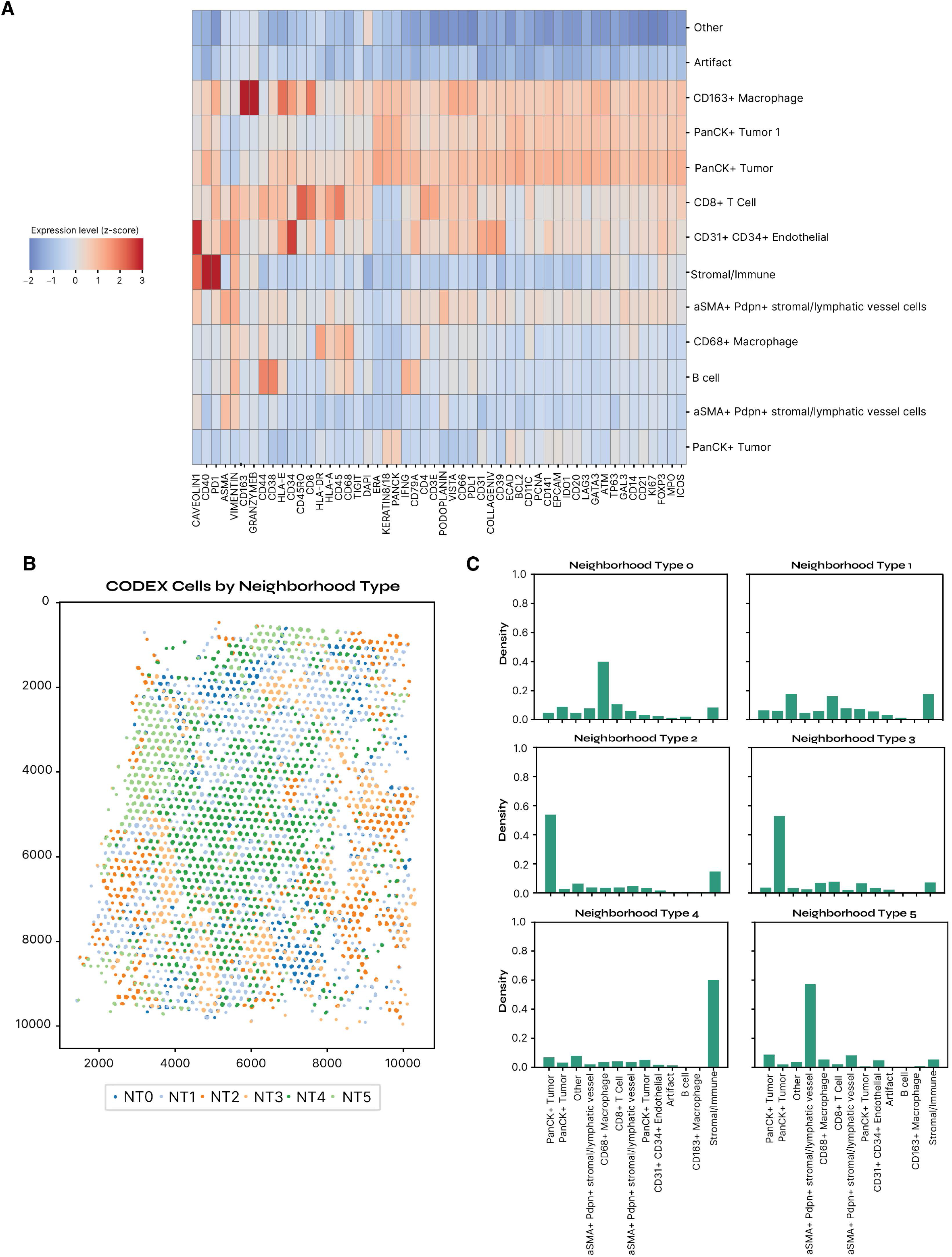
Defining cellular neighborhoods with CODEX data. **(a)** CODEX marker expression by celltype. Heatmap indicates the relative expression (color) of each biomarker (columns) in each cell type (rows). **(b)** Retained subset of CODEX cells (same as green points in Figure 3d) here colored by the Neighborhood Type to which they belong. **(c)** Cell type composition for each NT. Density (y-axis) of each cell type (x-axis) is shown for each NT (subplots).

