## Supplemental Table 1 for "emObject: domain specific data abstraction for spatial omics"

### Supplemental Table 1: Marker genes of Visium clusters

| V0 | V1 | V1 | V3 | V4 | V5 | V6 |
|---|---|---|---|---|---|---|
| ESR1 | FN1 | TCEAL4 | IGHA1 | IGKC | XBP1 | TMSB4X |
| BTG2 | TIMP3 | NPY1R | JCHAIN | IGHG1 | GJA1 | CD74 |
| MUC1 | ACTA2 | MUC1 | IGKC | IGHG2 | CYP4F8 | B2M |
| POSTN | MYL9 | SLC39A6 | CYP4F8 | IGLC1 | AZGP1 | HLA-DRA |
| AGR2 | DCN | SHROOM1 | IGLC1 | JCHAIN | PIP | FTL |
| STC2 | TAGLN | STC2 | IGHM | IGHG3 | CFD | PSAP |
| CITED4 | COL1A2 | SUSD3 | IGFBP3 | IGHM | MUC1 | HLA-DPB1 |
| NPY1R | LUM | TFF3 | AEBP1 | IGHA1 | SLC39A6 | LYZ |
| GFRA1 | THBS2 | CBLN2 | GJA1 | IGFBP7 | GLUL | APOE |
| EVL | AEBP1 | KRT18 | TIMP3 | VIM | MLPH | HLA-DPA1 |
| EXOC2 | ACTG2 | SERPINA3 | S100A6 | B2M | SERPINA3 | HLA-E |
| KRT8 | COL1A1 | CA2 | SHISA2 | C3 | SLC40A1 | TYROBP |
| CBLN2 | CCN2 | ESR1 | DCN | GSN | STC2 | LAPTM5 |
| CRACR2B | SPARC | AZGP1 | SEC14L2 | BGN | KRT18 | C3 |
| GATA3 | FLNA | TFF1 | SFRP2 | C1R | IL6ST | C1QB |
| PAQR4 | COL10A1 | CYP4Z1 | XBP1 | HLA-DRA | SEC14L2 | HLA-DQA1 |
| INPP4B | ACTB | UQCRQ | KLK11 | A2M | NFKBIZ | CTSB |
| CLEC3A | MMP2 | GFRA1 | SLC4A10 | IFITM3 | SPINT2 | IFI30 |
| NCAM2 | MMP14 | COX6C | IGHG3 | TMSB4X | MIF | APOC1 |
| BAMBI | MXRA8 | AGR2 | GAS6 | AQP1 | ASAH1 | HLA-DMB |
| BICDL2 | COL12A1 | ELOVL5 | TPSB2 | CCDC80 | CA2 | IL32 |
| SOWAHA | COL11A1 | ZNF552 | CLU | SOD3 | PPDPF | CCL5 |
| WFDC2 | DPYSL3 | GATA3 | PIP | FLNA | ELOVL5 | CTSD |
| VSIG2 | LOXL1 | CXCL14 | IGHG2 | SFRP4 | SCD | TXNIP |
| CAPN13 | POSTN | ELP2 | RIN2 | APOD | CFB | HLA-F |
| CDC20B | FBN1 | MAL2 | CDSN | HLA-DPA1 | ESR1 | TRAC |
| LRRC37A3 | SULF1 | GLUL | IGKV4-1 | C1QC | KRT8 | ITGB2 |
| BCL2 | RARRES2 | NAT1 | MFGE8 | HLA-DPB1 | VSTM2A | COTL1 |
| FOXA1 | VCAN | TBC1D9 | SLC40A1 | CD74 | CYP4Z1 | GPNMB |
| PPA2 | TIMP2 | VAV3 | COL10A1 | ID3 | SHISA2 | C1QC |
